# Templated paired helical filament formation and tau fold-dependent inclusion morphologies in primary neurons

**DOI:** 10.64898/2026.09.30.755038

**Authors:** Melissa Huang, Sofia Lövestam, Kimberly Ai Xian Cheam, Bernardino Ghetti, William A. McEwan, Sjors H. W. Scheres, Michel Goedert

**Affiliations:** MRC Laboratory of Molecular Biology, Cambridge, UK; UK Dementia Research Institute at the University of Cambridge, Department of Clinical Neurosciences, Cambridge, UK; Department of Pathology and Laboratory Medicine, Indiana University School of Medicine, Indianapolis, USA

**Keywords:** tau, amyloid, Alzheimer’s disease, neurodegeneration

## Abstract

Neurodegenerative tauopathies are characterized by the abnormal assembly of tau into amyloid filaments with distinct, disease-specific folds. These filaments are proposed to spread in a prion-like manner, thus underlying the progression of pathology in diseased brains. A prediction of this hypothesis is that tau filaments will induce progeny filaments identical to the input seed material, yet cellular models of templated tau assembly have failed to fully recapitulate brain-derived filament structures. Here, we show that mixed primary mouse cortical neurons overexpressing full-length human tau via adeno-associated virus transduction propagate Alzheimer’s disease paired helical filaments upon exposure to human brain-derived seeds. Using minimal tags and core mutations, we were able to unambiguously differentiate induced aggregates from the original seeds. Super-resolution microscopy and live-cell imaging revealed progressive inclusion formation in neurons. Moreover, seeding primary neurons with brain extracts from individuals with Pick’s disease, corticobasal degeneration and progressive supranuclear palsy induced the formation of morphologically distinct inclusions. Together, these findings establish that primary neurons provide an environment for the faithful propagation of paired helical filaments and show that tau filament folds dictate higher-order inclusion morphologies within neurons.

**Significance Statement:** Neurodegenerative tauopathies are characterized by the assembly of tau protein into disease-specific amyloid folds. While prion-like propagation is hypothesized to drive disease progression, cell models have previously failed to fully replicate human-brain derived filament structures. Here we show that primary mouse neurons support the templated propagation of human Alzheimer’s disease paired helical filaments. By incorporating a disease-causing mutation (V337M) into the amyloid core, we unambiguously resolved cryo-EM structures of the induced assemblies. Moreover, by using super-resolution light microscopy, we show that seeds from tauopathies induced the formation of morphologically distinct intraneuronal inclusions. These findings describe a system for dissecting the cellular factors that govern tau fold propagation and for understanding how distinct filament structures dictate phenotypic differences across tauopathies.

## Introduction

Neurodegenerative tauopathies, including Alzheimer’s disease (AD), Pick’s disease (PiD), progressive supranuclear palsy (PSP), and corticobasal degeneration (CBD) are characterized by the assembly of microtubule-associated protein tau into amyloid filaments (*1*). Mutations in *MAPT*, the gene encoding tau protein, cause dominantly inherited forms of frontotemporal dementias with abundant tau inclusions, consistent with tau filament formation causing neurodegeneration (*2–4*).

Tau is an intrinsically disordered protein that is expressed as six different isoforms in the adult human brain (*5*). These isoforms are produced by alternative mRNA splicing and differ by the number of microtubule-binding repeats (3R or 4R) and the number of N-terminal inserts (0N, 1N, or 2N). Tau filaments are made of a structured core and an unstructured fuzzy coat, with each tauopathy having a characteristic fold (*6*). Cryo-EM has shown that the minimal tau filament core consists of R3, R4 and 10-12 amino acids after R4. Tauopathies fall into three groups, based on the tau isoforms present in the filaments. In AD, all six tau isoforms (0N3R, 1N3R, 2N3R, 0N4R, 1N4R, 2N4R) assemble into paired helical and straight filaments (PHFs and SFs) (*7, 8*). PSP and CBD each have distinct filament structures that are made of 4R tau isoforms (0N4R, 1N4R, 2N4R) (*9, 10*), whereas 3R tau isoforms (0N3R, 1N3R, 2N3R) are found in the filaments of PiD (*11–13*). Tauopathies differ from each other not only at the molecular level, but also by their cell type vulnerability, neuroanatomical progression and clinical manifestation. Notably, tau inclusions are morphologically distinct in different tauopathies, which may indicate differences in inter-filament packing and organization within the cell (*14*). The development of cellular model systems that recapitulate the tau filament structures found in disease is therefore essential for understanding the molecular mechanisms of tauopathies.

Exogenously applied tau filaments can induce the assembly of intracellular monomeric tau into filaments. This results in the propagation of tau filaments in a prion-like manner and is believed to underlie the progression of disease and the spreading of tau pathology along neuroanatomical connections (*15*). When tau filaments are injected into mouse brains, they are taken up at the injection sites, where they induce further tau assembly, after which pathology spreads to distal brain regions, suggesting cell-to-cell propagation of tau filaments (*16–21*). Moreover, the injection of tau filaments extracted from the brains of individuals with different tauopathies into the brains of transgenic mice results in tau pathology with characteristics reminiscent of the human diseases, both in terms of cell-type specificities and morphologies (*17, 18, 22*).

The seeded assembly of monomeric tau into filaments has been demonstrated in cell culture systems. The addition of brain-derived tau seeds to HEK293 cells overexpressing wildtype or mutant tau has resulted in the robust assembly of monomeric tau into filaments (*23–28*). However, where investigated, their cryo-EM structures differed from those of the seeds. Specifically, undifferentiated SH-SY5Y cells that transiently expressed full-length wild-type human tau propagated only single protofilaments with the Alzheimer fold (*27*), and HEK293 cells stably expressing residues 297-391 of human tau propagated two protofilaments with the Alzheimer fold, but with different inter-protofilament packing from that of PHFs (*25*). Thus, while mounting evidence supports faithful prion-like replication of tau filaments, existing cellular models do not support the propagation of disease-relevant tau structures.

Here we report that mixed primary mouse cortical cultures overexpressing HA-tagged full-length human tau in neurons through AAV transduction form abundant HA-positive tau filaments upon exposure to seeds from the brains of individuals with AD. By cryo-EM, the templated filaments were identical to PHFs from AD brains, providing evidence that a murine neuronal environment can be compatible with the formation of human brain-derived tau filament structures. Moreover, super-resolution light microscopy showed that seeded aggregation with AD, PiD, PSP and CBD-derived tau seeds in 3R or 4R tau-expressing neurons led to seeded assemblies with distinct morphological characteristics.

## Results

### Seeded aggregation of tau inclusions in transduced primary mouse cortical neurons

To study the seeded aggregation of tau in neurons, we developed a system using human brain-derived tau seeds and adeno-associated virus (AAV)-mediated neuronal overexpression of hemagglutinin (HA)-tagged full-length human tau protein (**Figure 1A**). The HA-tag enables the detection of seeded tau aggregates rather than that of the added seeds. Mixed primary cultures contained neurons (30±8%, n = 4), astrocytes (47±16%, n = 4), and microglia (5±4%, n = 4) at DIV14, which represents a cellular environment that is more like that of the brain than are monocultures. We used chimeric particles of an AAV to deliver a construct encoding HA-tagged 0N3R or 0N4R human tau under the control of a neuron-specific promoter at DIV0 **(Figure 1A**). This resulted in high transduction efficiency (78±6%, n = 3) and human tau expression specifically in neurons (**Figure S1)**. The expression pattern of tau was diffuse and localized to both somatic and dendritic compartments (**Figure 1A, Figure S1**). The addition of AD brain-derived tau seeds at DIV3 to primary cultures overexpressing either HA-0N3R or HA-0N4R tau led to the formation of perikaryal and neuritic inclusions (**Figure 1B**). Tau inclusions were recognized by the HA tag and co-localized with phosphorylation-dependent anti-tau antibody AT100 (pT212/S214/T217) following methanol fixation (**Figure 1B**). Sarkosyl-insoluble tau extracted from neuronal cultures at DIV14 was detected by immunoblotting with antibodies against the C-terminus of tau (BR134) and the HA tag. (**Figure 1C**). Immuno-EM of the sarkosyl-insoluble extracts confirmed the presence of HA-labelled tau filaments (**Figure 1D**). Deletion of amino acids 306-311, which are required for tau filament formation (*29–32*), prevented the formation of HA-tau inclusions and sarkosyl-insoluble tau (**Figure 1B, C**). Consistent with previous reports (*19, 22*), a few HA-negative neuritic inclusions were observed with anti-tau antibody AT100, indicating the assembly of endogenous mouse tau. However, there was insufficient mouse tau in the sarkosyl-insoluble fractions to be detected by immunoblotting (**Figure 1C**). Thus, overexpression of human tau was required for the robust seeded aggregation of tau in primary neurons.

**Figure 1.**
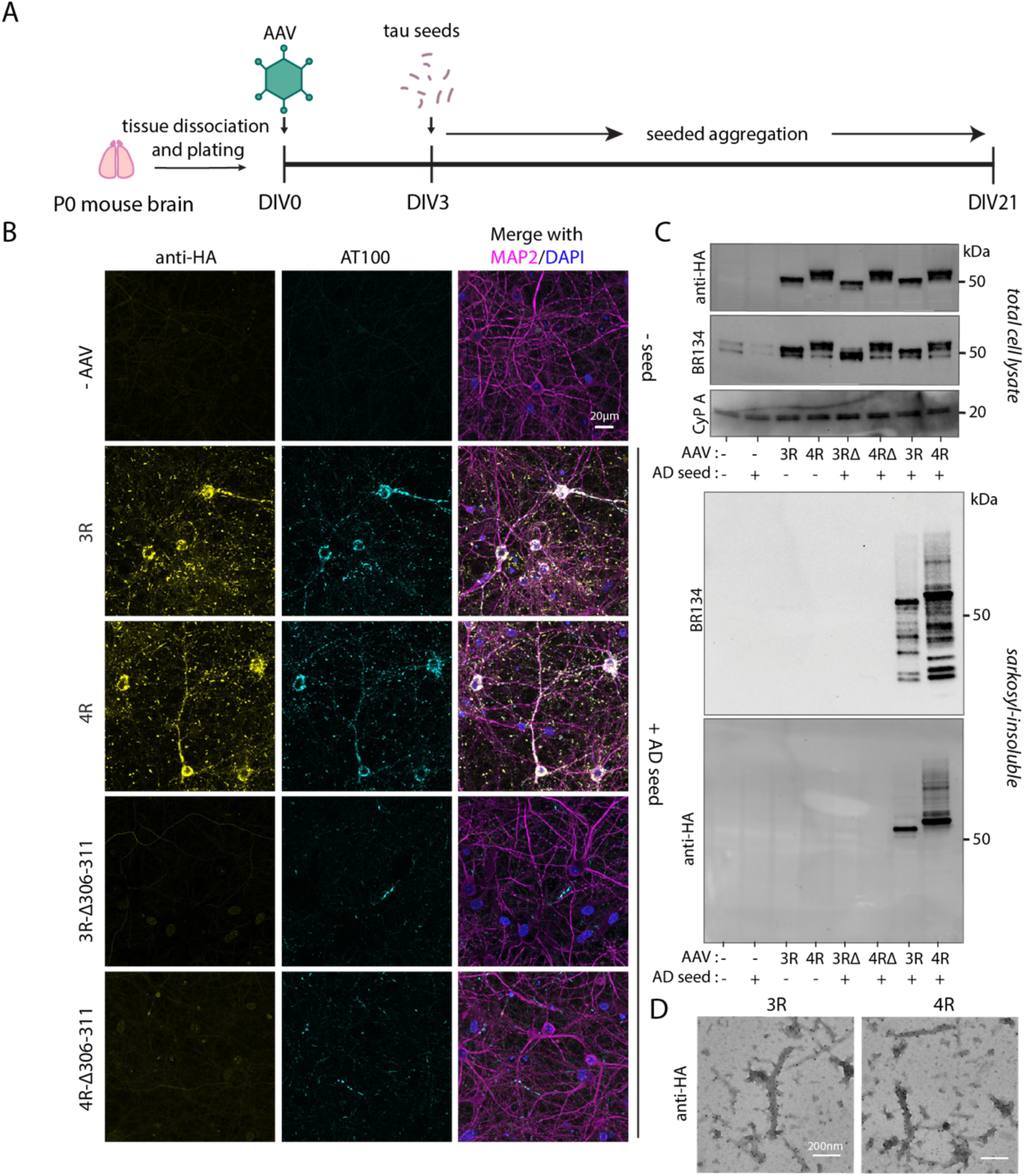
Seeded assembly of tau in AAV-transduced primary neurons. A) Schematic depicting the transduction and seeded aggregation of tau in primary mouse cortical neurons. B) Representative immunofluorescence images of primary cultures at DIV21 which were not transduced (-AAV) or transduced with chimeric particles of AAV (AAV1/2) to express tau under the neuronal-specific human synapsin-1 promoter (hSyn) with AAV1/2-hSyn-HA-0N3R (3R), AAV1/2-hSyn-HA-0N4R (4R), AAV1/2-hSyn-HA-0N3R-Δ306-311 (3R-Δ306-311), or AAV1/2-hSyn-HA-0N4R-Δ306-311 (4R-Δ306-311). Deletion of amino acids 306-311 prevented the formation of filaments. Cultures were untreated (-seed) or treated with AD seeds. Methanol-resistant anti-HA+ and AT100+ aggregates in MAP2+ neurons appeared in 3R or 4R tau-expressing neurons treated with AD seeds. Small amounts of neuritic, AT100+ staining, but no perikaryal anti-HA+ aggregates, were visible in primary neurons expressing 3R-Δ306-311 or 4R-Δ306-311 tau and treated with AD seeds. Cells were fixed with cold methanol. C) Immunoblots with anti-HA and BR134 (tau C-term) of total cell lysates and sarkosyl-insoluble tau extracted from primary cultures expressing 3R, 4R, 3R-Δ306-311 (3RΔ), or 4R-Δ306-311(4RΔ) at DIV21. D) Immunoelectron micrographs depicting the sarkosyl-insoluble extracts using anti-HA antibodies showing immunogold labelled filaments.

We next sought to establish the timeframe in which seeded aggregation occurs in primary neurons. We monitored seeded tau aggregation over 18 days (DIV3-21) following the addition of AD seeds. In untransduced cultures, the AT100 inclusions appeared in neurites at DIV10 and accumulated over time (**Figure 2A**). In cultures overexpressing HA-0N3R tau, the first inclusions were also observed at DIV10; however, they were more numerous and were present in both cell bodies and neurites (**Figure 2A**). This was accompanied by the presence of sarkosyl-insoluble tau (**Figure 2B**). This AAV-based expression system facilitates the genetic manipulation of tau constructs. To monitor tau aggregation in living neurons, we fused the fluorescent protein dendra-2 to the N-terminus of tau. Live-cell imaging over three weeks showed the progressive appearance of bright inclusions, a proportion of which moved along neuronal processes (**Movie S1**). With confocal live-cell imaging, we observed multiple independent fluorescent inclusions forming in neuronal cell bodies, where their sizes and intensities increased over time (**Movie S2**). Taken together, these experiments show that seeded aggregation can be monitored using N-terminal protein fusion tags and that it increases over time.

**Figure 2.**
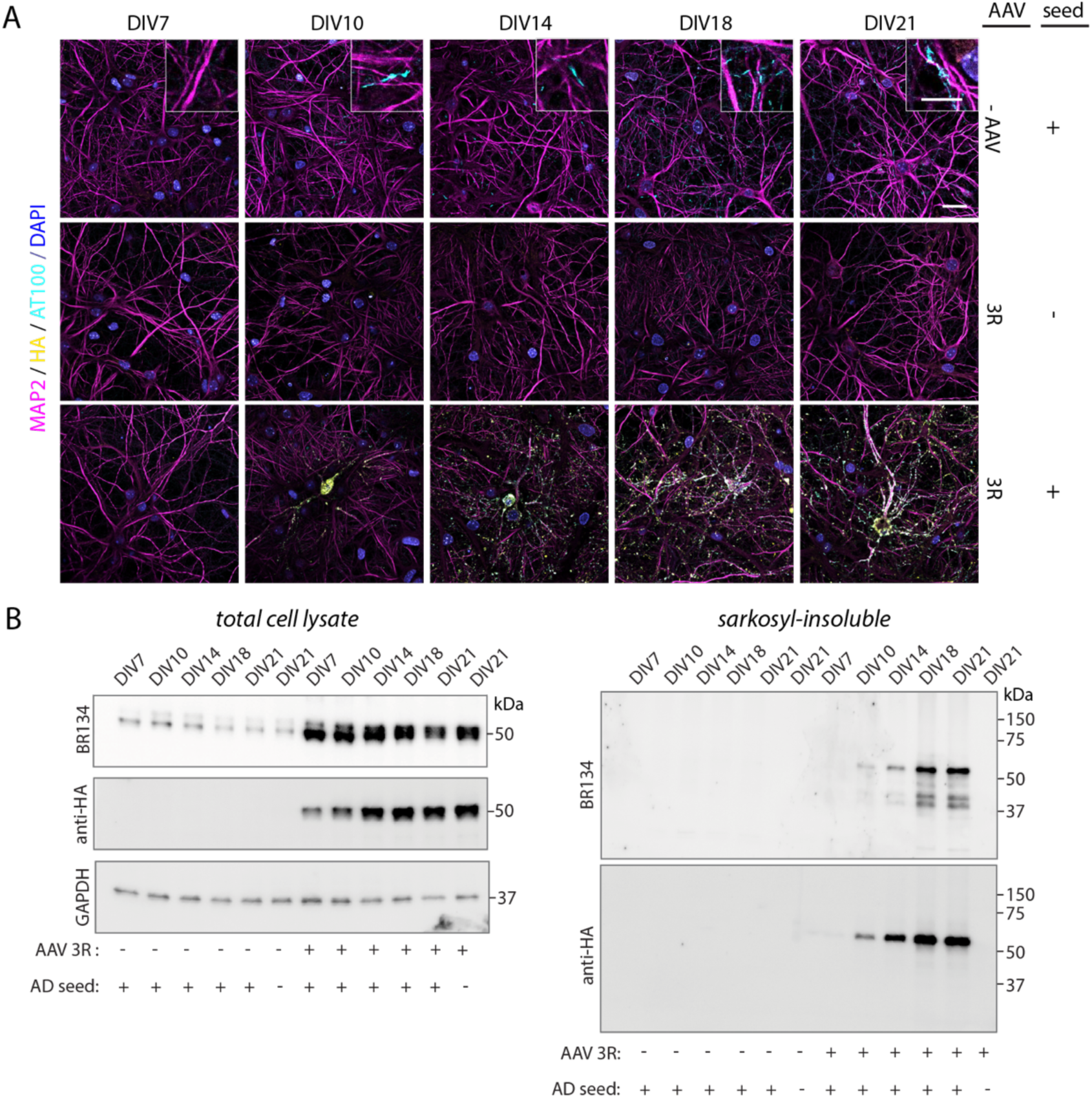
Progression of tau inclusions in AD-seeded primary neurons. A Representative immunofluorescence images of untransduced (-AAV) (top) and HA-0N3R tau-expressing (middle and bottom) primary cultures at DIV7, DIV10, DIV14, DIV18, and DIV21 following treatment with AD seeds, using anti-HA and AT100 antibodies. B) Immunoblots of total cell lysates and sarkosyl-insoluble fractions of untransduced (-) or AAV1/2-hSyn-HA-0N3R tau-transduced (+) neurons at DIV7-21, +/- AD seeds using BR134 and anti-HA antibodies.

## Cryo-EM structures of seeded tau filaments

We next sought to determine the cryo-EM structures of tau filaments extracted at DIV21 following the addition of AD seeds (0.2 mg of brain tissue per cm^2^ of cell culture surface area) at DIV3. The HA tag is located in the unstructured fuzzy coat, meaning that it can be accessible by immuno-EM, but that it is not resolved by cryo-EM. We therefore established an alternative method to unambiguously demonstrate that we were solving the cryo-EM structures of the templated filaments rather than those of the seeds. This method consisted in the overexpression of HA-0N3R tau with the human *MAPT* mutation V337M (*2*). This mutation resides within the core region of tau filaments and gives rise to inherited frontotemporal dementia and PHFs that are identical to those of AD (*33*). We used this property to enable positive identification of the seeded filaments. Thus, high-resolution structures could discriminate between seeds and seeded assemblies, since the seeds from the brains of AD patients were V337 and the templated filaments were M337. The addition of AD seeds to neuronal cultures overexpressing HA-0N3R-M337 tau led to the formation of HA-positive neuronal inclusions that co-localized with the pentameric oligothiophene amyloid dye (pFTAA) and anti-tau antibody AT100 (**Figure S2)**. Immuno-EM of the sarkosyl-insoluble extracts confirmed the presence of HA-labelled tau filaments (**Figure S2**).

Cryo-EM analysis showed that the majority (82%) of tau filaments extracted from primary neurons were PHFs and that the density at residue 337 was that of methionine (**Figure 3A**). We also observed density for residues 379-381 turning inwards, towards the tip of the protofilament. A similar density was observed for the singlets that resulted from seeding SH-SY5Y cells (*27*), but this density was not present in PHFs from AD brains (*7, 8*). There was also a minor population (18%) of single protofilaments with the Alzheimer tau fold (**Figure S3**). At a resolution of 2.9 Å, the side chain density at position 337 in the singlet map was less clear. However, rigid-body alignment of the atomic model of one of the protofilaments of the PHF with a methionine provided an excellent fit to the singlet map, including the density turning inwards, indicating that the singlets were also the products of seeded aggregation. We thus established that the filaments used for cryo-EM structure determination were the products of seeded aggregation and that they were predominantly PHFs.

**Figure 3:**
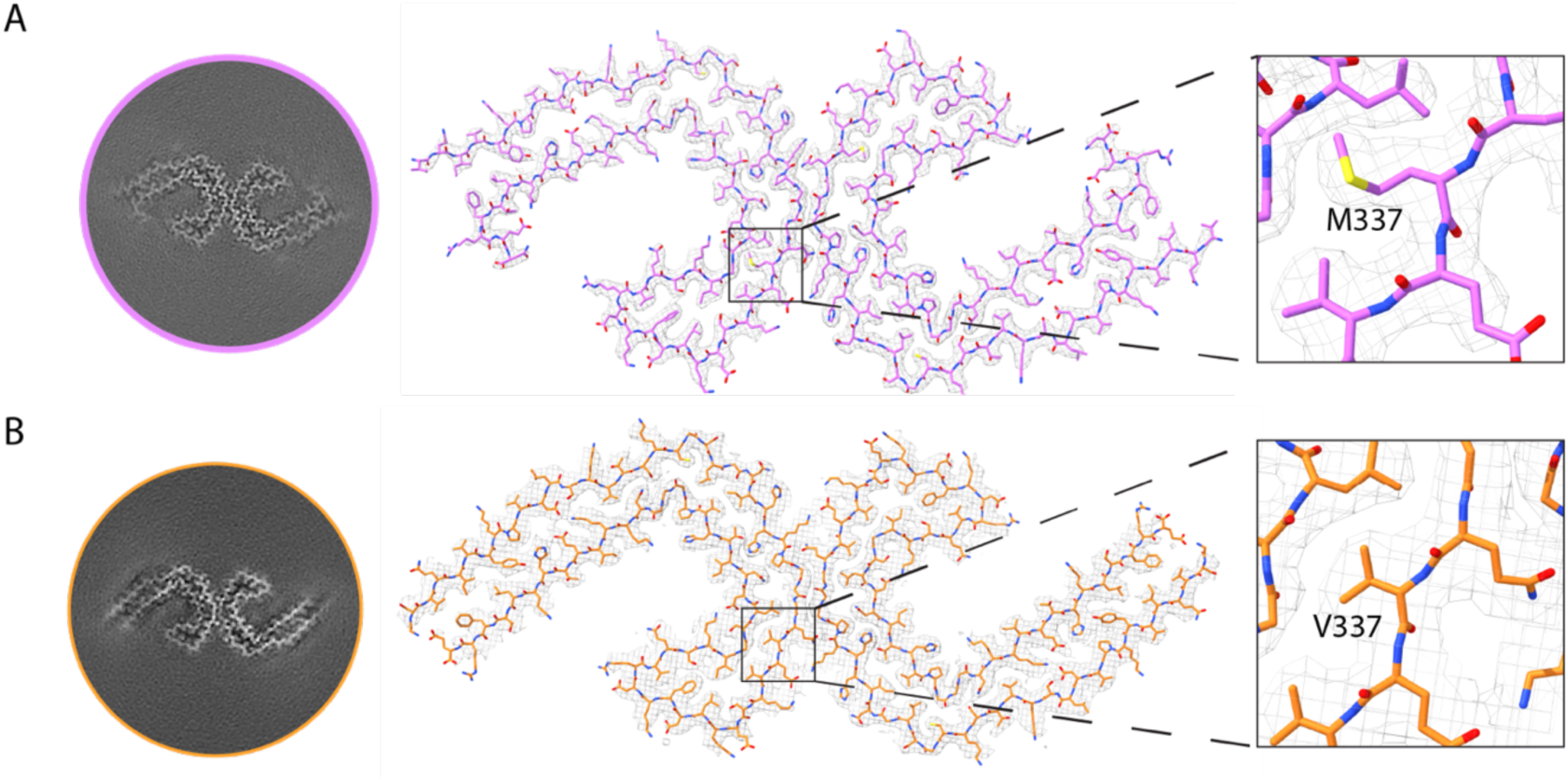
Cryo-EM densities and atomic models of PHFs derived from primary neurons. A) Cross-section of reconstructed density and cryo-EM map of filaments at DIV21 that were extracted from primary cultures expressing HA-0N3R-M337 tau and treated with AD seeds (0.2mg/cm^2^). The inset shows a methionine residue at position 337, proving that the tau filaments had formed by seeded aggregation. PHFs represented 82% of filaments and were solved to a resolution of 2.3 Å. The remainder (18%) were tau singlets with the Alzheimer fold. B) Cross-section of reconstructed density and cryo-EM map of filaments extracted from primary cultures expressing HA-0N3R-M337 tau and treated with AD seeds (8mg/cm^2^). The inset shows a valine residue at position 337, indicating that the original AD seeds had been re-extracted and their structures solved. PHFs represented 100% of filaments and were solved to a resolution of 2.8 Å.

When AD seeds were added in excess (8 mg/cm^2^), tau filaments were again predominantly PHFs. However, the density at residue 337 was that of valine and there was no density turning inwards for residues 379-381, indicating that we had solved the structures of the original seeds (**Figure 3B, Figure S3)**. Careful titration of seed concentrations is therefore essential for distinguishing between the templated assemblies and residual seeds.

### Isoform-specific seeding of 3R and 4R tauopathies in primary neurons

We next investigated whether this platform could also be applied to other tauopathies, specifically PiD, CBD and PSP. The AD fold excludes R1 and R2 of the microtubule-binding repeat domains and incorporates both 3R and 4R tau isoforms. In contrast, other tauopathies have filament structures that permit incorporation of only 3R (PiD) or 4R (CBD, PSP) tau isoforms. Seeded aggregation of tau in primary neurons using AD, PiD, CBD and PSP seeds exhibited isoform specificity (**Figure S4)**. AD seeds induced the formation of HA-tagged sarkosyl-insoluble tau in neurons expressing either HA-0N3R or HA-0N4R, whereas PiD seeds specifically induced HA-0N3R aggregation, and CBD and PSP seeds induced HA-0N4R aggregation.

Addition of AD or PiD seeds to neurons expressing HA-0N3R tau and AD, CBD or PSP seeds to neurons expressing HA-0N4R tau resulted in the formation of perikaryal and neuritic inclusions that were positive for HA and pFTAA (**Figure 4A**). We performed immuno-EM on the sarkosyl-insoluble fractions of these seeded cultures and observed HA-labelled filaments (**Figure 4B, C, Figure S4**). Together, this shows that tau seeds from different tauopathies can induce HA-tagged filamentous neuronal inclusions in an isoform-dependent manner.

**Figure 4:**
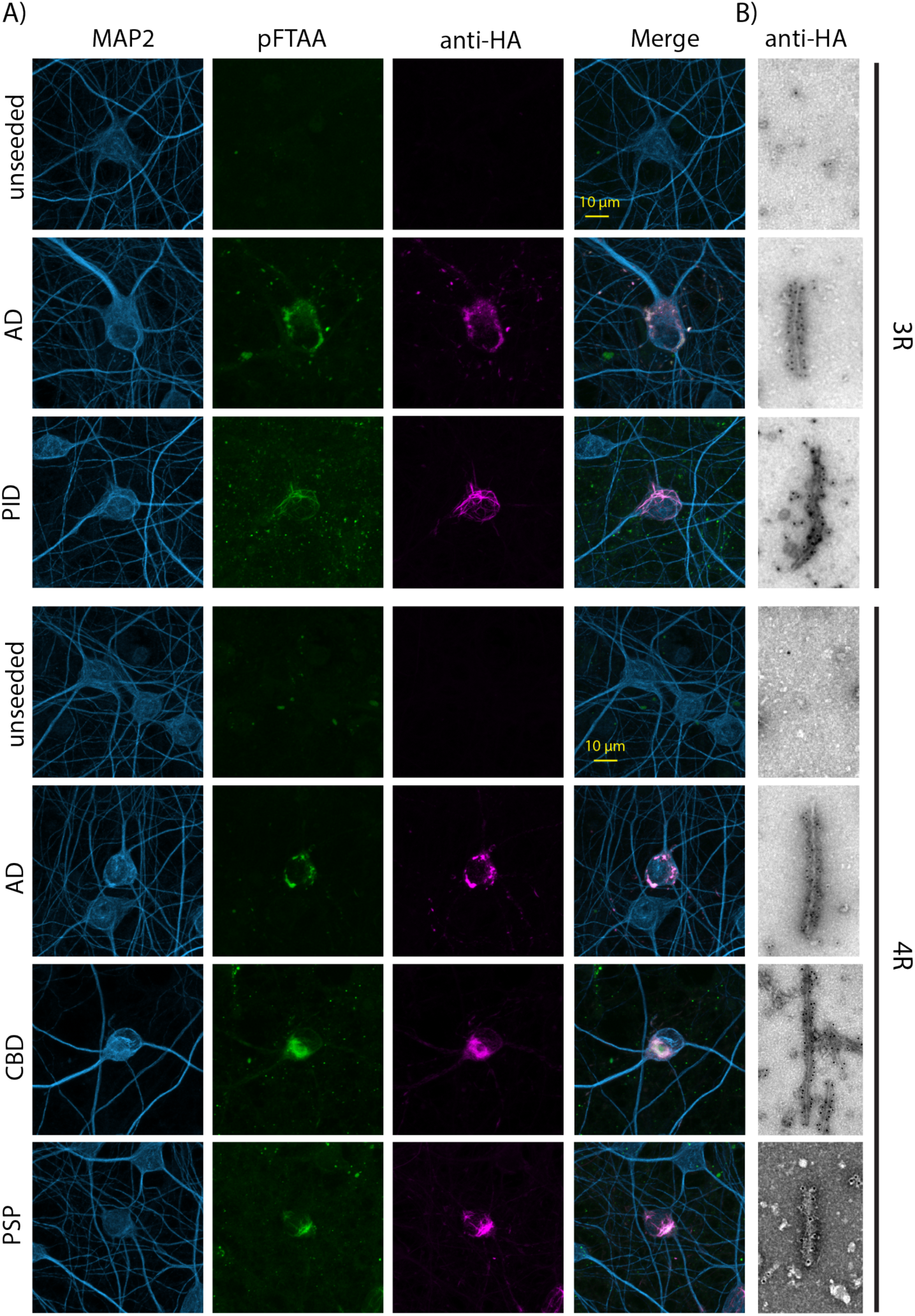
Seeded aggregation with AD, PiD, CBD and PSP-derived seeds in primary neurons. A) Representative immunofluorescence images of primary neurons at DIV21 overexpressing HA-0N3R tau (3R) or HA-0N4R tau (4R) (bottom panel) and treated with AD, PiD, CBD or PSP seeds. B) Immuno-EM of sarkosyl-insoluble fractions showing immunogold-labelled filaments with anti-HA antibody.

### Tau fold-dependent inclusion morphologies

Given that tau filaments have specific atomic structures in different tauopathies (*6*), we wondered whether there might be morphological differences in the seeded neuronal inclusions visible by stimulated emission depletion (STED) super-resolution light microscopy. PHFs have widths of 7nm at the narrowest and 15nm at the widest and can be several microns in length (*7*). We used a primary anti-HA antibody and STAR-RED-conjugated secondary antibody to label the inclusions following seeded aggregation, which would add up to 30nm to a labelled tau filament (*34*). In unseeded neurons, we observed diffuse, intraneuronal HA labelling. An AD-seeded HA-tau inclusion was resolved using STED imaging to full width at half-maximum (fwhm) of 71nm, whereas the width by diffraction-limited imaging was 210nm (**Figure S5**). Tau inclusions appeared filamentous in morphology, with an average length of 612nm±236.3 (n = 202), ranging from 137-1427nm. We performed a time-course analysis by fixing AD-, PiD-, CBD- and PSP-seeded neurons at DIV7, DIV10, DIV14, DIV18 and DIV21. The first filamentous inclusions appeared at DIV10 and increased in density until DIV14 (**Figure 5**). At DIV18, we also observed dense clusters of HA-tau inclusions, the extent of which increased by DIV21. At DIV18 and DIV21, we saw striking morphological differences between AD-, PiD-, CBD- and PSP-seeded cell body inclusions (**Figure 5, Figure S6**). Whereas AD-seeded inclusions formed dense clusters, we observed long and wide ribbon-like inclusions in PiD-seeded neurons. CBD- and PSP-seeded neurons had dense networks of long, thin filamentous inclusions, with occasional thicker fibers. Thus, mouse neurons support the propagation of tau pathology with morphological characteristics specified by the brain-derived seeds.

**Figure 5:**
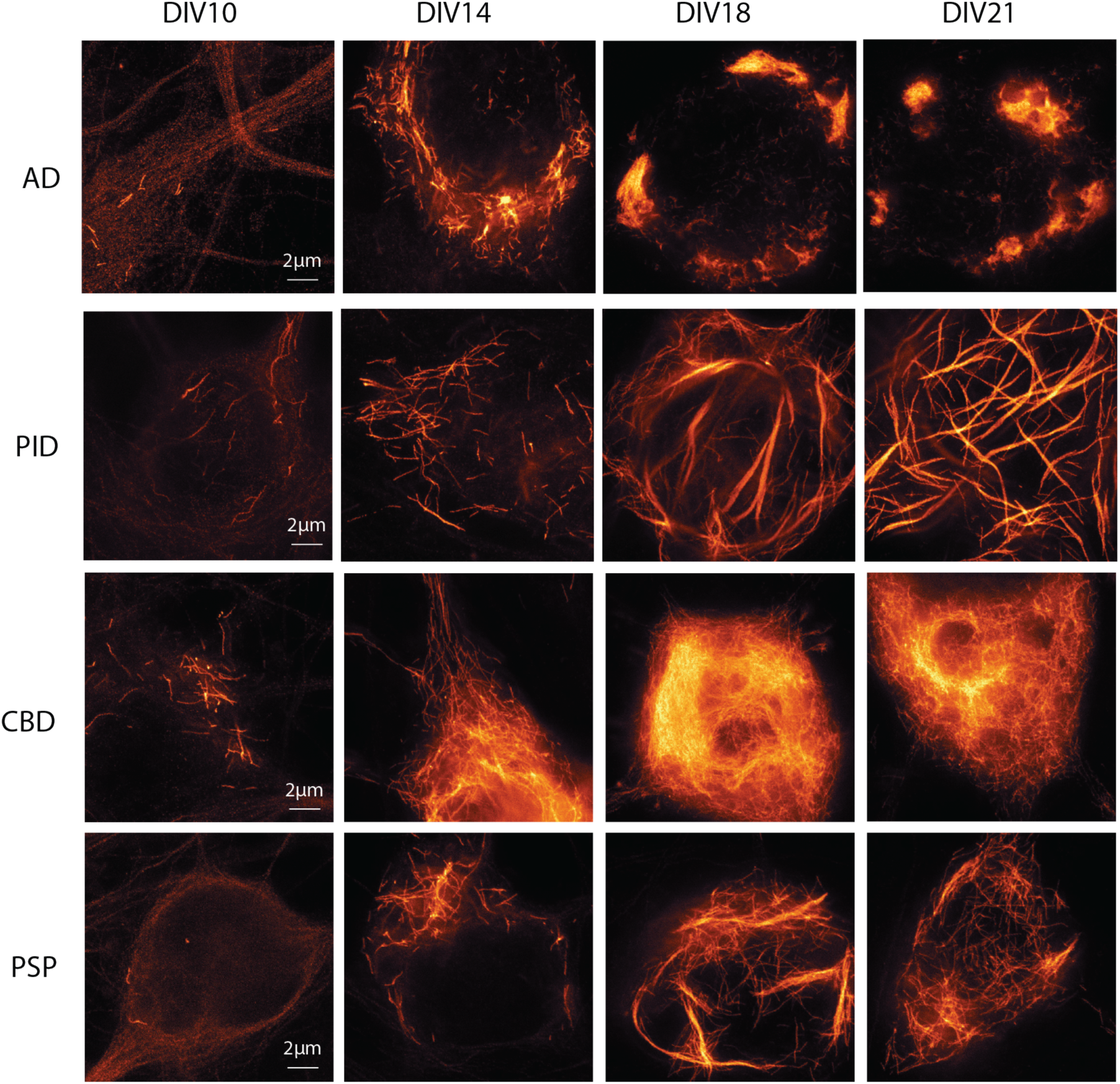
Tauopathy-derived seeds produce distinct intraneuronal inclusion morphologies. Representative STED images of primary neurons overexpressing HA-0N3R tau and seeded with AD or PiD tau seeds, or HA-0N4R tau and seeded with CBD or PSP tau seeds. Neurons were fixed with methanol and immunolabelled with an anti-HA antibody at DIV10, DIV14, DIV18 and DIV21.

## Discussion

Here we describe a neuronal culture system of seeded aggregation using AAV-mediated overexpression of full-length human tau protein. We demonstrate that brain-derived tau seeds from AD cases drove the robust seeded aggregation of tau into neuronal inclusions composed of PHFs. This is the first time that human tau was demonstrated to be experimentally templated into PHFs inside cultured neurons, in support of the prion-like tau spread. Moreover, these findings suggest that murine primary neurons can provide a cellular environment that is compatible with such propagation. The underlying mechanisms are not known, but they may be critical for understanding the spread of pathology within the brain. We identified a minority population of single protofilaments with the Alzheimer tau fold, like those found in undifferentiated SH-SY5Y cells (*27*). This suggests that differences in the duration of the seeding experiments or the cellular environments could contribute to the formation of PHFs.

We used an HA tag at the N-terminus and a *MAPT* mutation (V337M) in the core of PHFs to detect seeded tau aggregates. In biochemical experiments, including immunofluorescence, immunoblotting and immuno-EM, the tag can be used to discriminate between cellular tau and the seeds. However, for cryo-EM studies, where the N-terminal tag cannot be resolved, we introduced a mutation in the cores of the amyloid filaments to discriminate them from the added seeds. We used the inherited *MAPT* mutation encoding V337M tau, which gives rise to frontotemporal dementia and PHFs (*2, 33*), to show that we solved the cryo-EM structures of the seeded tau filaments, rather than those of the seeds. At high seed concentrations, it was possible to re-extract the seeds. Thus, careful consideration must be applied to the amount of tau seeds used. These findings underline the importance of using untransduced cells, in conjunction with negative-stain EM and immunoblotting, as controls to determine the concentrations at which seeds are not re-extracted.

A neuronal cell culture system that can recapitulate disease-relevant folds is essential for advancing our understanding of tau filament formation in neurodegenerative diseases. An advantage of using AAV-mediated transduction to drive tau overexpression is its versatility, which addresses bottlenecks inherent to transgenic mouse models and iPSC-derived neuronal cell lines. We demonstrate this versatility by monitoring seeded aggregation over time using either an HA-tag or a fluorescent dendra-2 tag. In future experiments, appending various effector tags, such as a HaloTag for single-particle tracking and TurboID for proximity labelling, will open ways for studying the dynamics of tau aggregation in living neurons and for identifying proteins that interact with tau during filament formation. These tools will be critical for the identification of chaperones and degradation machinery that may engage with disease-specific folds.

Viral delivery also permits the study of the effects of disease-causing and other *MAPT* mutations. Such experiments will enable dissection of the requirements for tau filament formation and tau fold-specific propagation in neurons. Moreover, while the current study used the hSyn neuron-specific promoter, the mixed primary cultures also contain significant populations of astrocytes, which may similarly be genetically manipulated by AAV transduction. Swapping hSyn for an astrocyte-specific promoter will enable this platform to be used for studying tauopathies with a substantial astrocytic component, such as CBD and PSP. The same framework could also be used to drive the expression of other proteins that assemble intracellularly in neurodegenerative diseases, such as α-synuclein, TDP-43 and TAF15.

Potential limitations of AAV-mediated overexpression must also be considered. Although AAV particles are less immunogenic than other viral vectors, high viral titres might still activate inflammatory responses. Moreover, driving high-level expression of human tau protein carries the risk of overexpression artifacts. We found that overexpression is necessary for robust seeding of tau within the time frame of primary neuronal cultures; it remains to be seen if it is possible to use lower virus titres.

Using STED imaging, we further show that AD-tau seeded neuronal inclusions were similar to those from AD brains (*35*). By examining different timepoints following seeded aggregation, we show a progression in morphology from discrete filamentous inclusions to dense clusters and bundles of tau inclusions. Further time-resolved super-resolution microscopy experiments will enable us to study the dynamics of tau aggregation and filament formation in a neuronal context. We also show, by STED imaging, that tau seeds derived from tauopathies with different filament structures induce morphologically distinct inclusions in neurons. Super-resolution microscopy has not been performed on PiD, CBD, or PSP brain tissues. It therefore remains to be seen if these morphological differences resemble those found in the human brain. These findings, however, suggest a direct link between the underlying conformations of the tau seeds and the morphologies of the neuronal inclusions. Future studies will determine the cryo-EM structures of tau filaments seeded with PiD, PSP, and CBD extracts.

In conclusion, we demonstrate by cryo-EM that PHFs form in a structurally faithful, prion-like manner following seeded aggregation in mouse neurons expressing human tau. Super-resolution microscopy has suggested that tau assemblies extracted from other tauopathy brains may also induce specific aggregation, with the propagation of isoform-selective and disease-specific morphologies, consistent with strain-like behavior. These results support intraneuronal seeded aggregation of human tau as a mechanism of pathology spread and establish robust protocols for studying the relationship between tau aggregation and its downstream effects.

## Materials and Methods

### Adeno-associated virus production

AAV Pro 293T (Takara) cells were maintained in Gibco DMEM + GlutaMAX + 10% FBS + 1% penicillin/streptomycin in a tissue culture incubator at 37°C with 5% CO_2_. The AAV transfer plasmid was prepared by cloning into pAAV-hSyn-eGFP (Addgene). AAV was produced by co-transfection of AAV Pro cells with Rep/Cap plasmid (pAAV2/1 and pAAV2/2, Addgene) and pAdDeltaF6 Helper plasmid (Addgene). Viral particles were purified by iodixanol gradient centrifugation, as described (*36*) and their purity was confirmed with SDS-PAGE and staining with Coomassie InstantBlue. Primary cultures were transduced on DIV0 by adding AAV into fresh maintenance media. The approximate viral titres were determined by qPCR using forward (GGAACCCCTAGTGATGGAGTT)) and reverse (CGGCCTCAGTGAGCGA) primers amplifying regions between the inverted terminal repeats (ITRs) and the optimal AAV multiplicity of infection (MOI) for expression with low toxicity was determined experimentally by serial dilutions of the virus onto primary cultures. Expression was quantified by fluorescence or immunofluorescence labelling, and cell viability was evaluated by PrestoBlue (A13261). Fluorescence signal from PrestoBlue reagent was read on a ClarioSTAR microplate reader (BMG Labtech) at 560/590 nm. The following viruses were used in this study at the given MOI [vector genomes (vg) / cell]: AAV1/2-hSyn-HA-0N3R (1,200 vg/cell) AAV1/2-hSyn-HA-0N4R (2,500 vg/cell), AAV1/2-hSyn-HA-0N3R-V337M (1,000 vg/cell), AAV1/2-hSyn-dendra2-0N3R (4,000 vg/cell).

### Mixed primary cortical cell cultures

Mixed primary cortical cultures were prepared as described (*37*). Briefly, hippocampal and cortical tissues from male P0 C57BL/6J mouse pups were sliced into ±8 pieces/brain and trypsinized (0.25%, Gibco) for 20min at 37°C, followed by the addition of 0.1% DNase I (Sigma-Aldrich) for 3 min. Trypsin was inactivated by washing in plating medium consisting of 10 ml Neurobasal plus (Thermo Fisher Scientific), 10% horse serum and heat-inactivated Glutamax (Invitrogen). The plating medium was then reduced to 2 ml, before titration with a fire-polished glass pipette (Merck Life Sciences). Cell viability and cell numbers were assessed using a Countess II cell counter slide. Cells were plated according to the number of live cells (96-well plate: 30,000 cells/well; 6-well plate: 600,000-800,000 cells/ well; 8-well chamber: 75,000 cells/well). Cultures were incubated at 37°C for 3-5 h, before the plating medium was removed and fresh medium was added. AAV was added during this medium change.

### Immunohistochemistry

Cells were fixed and permeabilized in 100% cold methanol for 5 min. Following fixation, methanol was removed by diluting 1:2 using ice-cold PBS, three times, before washing the cells in 100% PBS. Cells were blocked with PBS + 2% BSA and incubated in primary antibody at room temperature for 2 h or at 4 °C overnight. For expression analyses, cells were fixed at room temperature in 4% Image-iT paraformaldehyde (Thermo Scientific) for 10 min. Cells were permeabilized, blocked with PBS + 2% BSA + 0.1% Triton X-100 and incubated in primary antibody in blocking buffer at room temperature for 2 h or at 4 °C overnight. Nuclei were stained with DAPI or Hoechst 33342 (Thermo Scientific). The following primary antibodies were used: HT7 (Thermo Fisher, 1:1000), anti-HA-biotin (Roche, 1:500); AT100 (Invitrogen, 1:500), anti-MAP2 (Abcam, 1:10,000); anti-β-III-tubulin (Abcam, 1:200); anti-GFAP (Sigma-Aldrich, 1:500), anti-GFAP (Abcam, 1:500); anti-Iba-1 (Novus Biological, 1:200), anti-CypA (Biotechne, 1:1000). Alexa-fluor conjugated secondary antibodies (405, 488, 594, or 647) or streptavidin were used for labelling (Thermo Fisher, 1:1000). For STED microscopy, STAR RED and STAR ORANGE secondaries (Abberior, 1:200) were used, with ProlongGold anti-fade mounting medium (Thermo Fisher). STAR RED was used for the labelling of HA-tau aggregates. All primary antibodies for STED were used at a concentration of 1:200.

### Light microscopy

High content wide-field microscopy was performed using a Ti2-E High Content Microscope (Nikon) with a 10X lens. Images were analyzed using NIS-Elements software (Nikon) for analysis of percentage of cell types and transduction efficiency from multiple independent primary cultures.

Confocal images were acquired on a Stellaris Lighting Confocal Microscope (Leica) with a 63X lens, Zeiss 900 inverted microscope with a 40X lens, or a W1 Spinning Disk (Nikon), with a 60x lens. An Okolab chamber was used to maintain 37 °C and 5% CO_2_ with humidity. Long-term live cell imaging experiments were performed in an IncuCyte S3 Live-Cell system with a 10X lens, with 37 °C and 5% CO_2_ with humidity.

STED was performed on a Leica TCS SP8 STED 3X gated microscope (Leica Microsystems) with a 100x/1.4NA oil immersion objective. Images were acquired with a 7x zoom factor and 1024×1024 pixel size (16nm/pix). Samples were irradiated with a pulsed white light laser and depleted with a pulsed STED laser line at 775nm. Signals were detected using HyD photon counting. Cell bodies (n = 15) were imaged at each time point using seeds derived from two cases of each tauopathy (AD, PiD, CBD, and PSP). Discrete filamentous inclusions were measured manually in ImageJ.

### Sarkosyl extraction of tau filaments from human brains

Sarkosyl-insoluble tau was extracted, as described (*38*). Briefly, frozen brain tissues (1.2-2.4g) were homogenized in 10X (v/w) extraction buffer containing 800mM NaCl, 10mM Tris-HCl pH 7.4, 2.5mM EDTA, 10% sucrose, 2% sarkosyl with Pierce Protease and Phosphatase Inhibitor Mini Tablets (Thermo Scientific). With the exception of AD case 2, all tauopathy cases and their tau filament structures have been described (*9, 10, 12, 39*) (Table 1). AD case 2 is a female who died aged 87 with a clinicopathological diagnosis of AD (Table 1). By cryo-EM, tau filaments were PHFs (Figure S3). Tissue homogenates were incubated at room temperature for 1 h, followed by a 20 min centrifugation at 13,000g. The supernatants were transferred to 1.5mL ultracentrifuge tubes, then spun at 124,500g for 1 h at room temperature. The pellets were resuspended in 1mL extraction buffer and re-ultracentrifuged. The resulting pellets were resuspended in 1mL 50mM Tris-HCl, 150mM NaCl and ultracentrifuged, as before. The pellets were resuspended in 250μL/g starting weight and sonicated with a metal probe (Misonix Microson Ultrasonic Cell Disruptor, setting 3) for 15 s. Tau seeds were aliquoted in Protein Lo-Bind Eppendorf tubes and frozen at -20°C until use. Seeds (0.1-8.0mg of brain, extracted, per cm^2^ of cell culture vessel) were added to primary mixed cortical cultures at DIV3 by diluting into media. The optimal concentrations of tau seeds were determined experimentally with serial dilutions and depended on the yield of filaments per brain extract.

**Table 1.** Brain tissue samples of tauopathy disease cases.

|  | **Cortical regions** | **Ages at death** | **Gender** | **Tau filament structures** |
| --- | --- | --- | --- | --- |
| **AD case 1** | Frontal | 81 | Female | (39) |
| **AD case 2** | Frontal | 87 | Female | Figure S3 |
| **PID (ΔK281) case 1** | Frontal | 54 | Male | (12) |
| **PID (ΔK281) case 2** | Frontal | 53 | Female | (12) |
| **CBD case 1** | Frontal | 52 | Male | (10) |
| **CBD case 2** | Parietal | 64 | Male | (10) |
| **PSP case 1** | Parietal | 59 | Male | (9) |
| **PSP case 2** | Frontal | 62 | Male | (9) |

### Sarkosyl extraction of tau filaments from seeded mixed primary cortical cells

Media were removed by aspiration and the cells washed 2X with PBS. They were then lysed in 250 μL/well extraction buffer [800mM NaCl, 25 mM Tris-HCl pH 7.4, 5mM MgCl_2_, 15% sucrose, 1% sarkosyl + Protease and Phosphatase inhibitor tablets (Pierce)] and collected by pipetting up and down. This was followed by a 1 s sonication with a metal probe sonicator (Misonix Microson Ultrasonic Cell Disruptor, setting 2), before incubation with NEBEXpress Salt Active Nuclease (M0764S) at 37 °C for 30 min with shaking (400 rpm). The lysates were cleared by a 5 min centrifugation at 5,000 g, before spinning the supernatants at 124,500 g at room temperature for 1 h. The pellets were washed 2x in TBS (150 mM NaCl, 50 mM Tris HCl pH 7.4) before resuspension in TBS (5 μl/well).

### Biochemical and ultrastructural analyses of tau filaments

Tau filaments extracted from primary cultures were diluted 10-fold and deposited on glow-discharged 400-mesh formvar/carbon film-coated copper grids (model CF400-Cu, EM Sciences) for 40 s. Grids were blocked for 10 min in 0.1% cold water fish skin gelatin in PBS before a 1 h incubation with primary antibodies diluted in blocking buffer. Grids were washed with blocking buffer and incubated with gold-conjugated anti-mouse IgG (Sigma-Aldrich) or anti-rabbit (Sigma-Aldrich) secondary antibody for 1 h. Following washing, the grids were stained with 4 μl of 2% uranyl acetate for 90 s. Images were acquired using a transmission electron microscope (Tecnai G2 Spirit) at 120 kV. An anti-HA antibody (1:50, Sigma-Aldrich) was used for immuno-EM.

### Electron cryo-microscopy

Sarkosyl-insoluble extracts were prepared from approximately 2.1*10^6 cells. They were applied to glow-discharged holey carbon grids (Quantifoil Au R1.2/1.3, 400 mesh) and plunge-frozen in liquid ethane using Vitrobot Mark IV (Thermo Fisher Scientific) at 95% humidity and 4°C. Cryo-EM images were acquired on a Titan Krios electron microscopy operated at 300 kV with a Falcon 4i direct electron detector (Thermo Fisher Scientific). For the dataset with a methionine at residue 337, we used a Selectris-X energy filter with a slit width of 10 eV. Images were recorded at a dose of 40 e^-^/Å^2^ using EPU software (Thermo Fisher Scientific). See **Table S1** for further details.

### Image processing

Datasets were processed in RELION using helical reconstruction for amyloid filaments (*40–42*). Movie frames were gain-corrected, aligned, and dose-weighted using RELION’s implementation of the motion correction program. Contrast transfer function (CTF) parameters were estimated using CTFFIND-4.1 (*43*). Filaments were picked manually or automatically (*44*) and segments were extracted with a box size of 1024 pixels, downscaled to 128 pixels. Reference-free 2D classification was performed to remove poor-quality particles. Selected class averages were re-extracted using a box size of 256 pixels. *Relion_helix_inimodel2d* was used to generate initial models *de novo* from 2D class averages based on filament crossover distances. 3D refinement was performed in RELION, with further optimization of helical twist and rise by local searches. Bayesian polishing and CTF refinement were then performed to increase resolution (*45–47*). Resolutions were determined based on Fourier shell correlations (FSC) between two independently refined half-maps at the 0.143 criterion (**Figure S7**).

### Model building and refinement

Atomic models were built using Model Angelo (*48*) or previously published structures as references (6HRE). Model refinements were performed using Servalcat, Coot, and Isolde (*49–51*). Models were validated with MolProbity (*52*). Figures were prepared with ChimeraX (*53*).

## Acknowledgments

We are grateful to the individuals and their families for donating brain tissues. We thank T. Darling, I. Clayson and J. Grimmett for help with high-performance computing, the EM facility of the Medical Research Council (MRC) Laboratory of Molecular Biology for help with cryo-EM data acquisition and the staff at ARES and the Biological Services Group for their help with animal husbandry and the collection of animal tissues. We thank N. Barry and the Light Microscopy facility of the MRC Laboratory of Molecular Biology for their help with STED microscopy.

## Data availability

Atomic coordinates for the seeded paired helical filaments with a methionine at position 337, the singlets with a methionine at position 337 and of the seeds with a valine at position 337 have been deposited at the Protein Data Bank (accession codes 33WM, 33WP and 34RF, respectively); corresponding electron microscopy maps have been deposited at Electron Microscopy Data Bank (accession codes EMD-59639, EMD-59642 and EMD-100131 respectively). All other materials are available upon request. Requests for materials should be addressed to W.A.M., S.H.W.S. and M.G.

## Author Contributions

M.H. and K.A.X.C. performed experiments, M.H. and S.L. collected cryo-EM data., B.G. identified Alzheimer’s disease case 2 and performed neuropathology. M.H., S.H.W.S, and M.G. wrote the manuscript. W.A.M., S.H.W.S, and M.G. supervised the project.

