## Supplementary information for "Templated paired helical filament formation and tau fold-dependent inclusion morphologies in primary neurons"

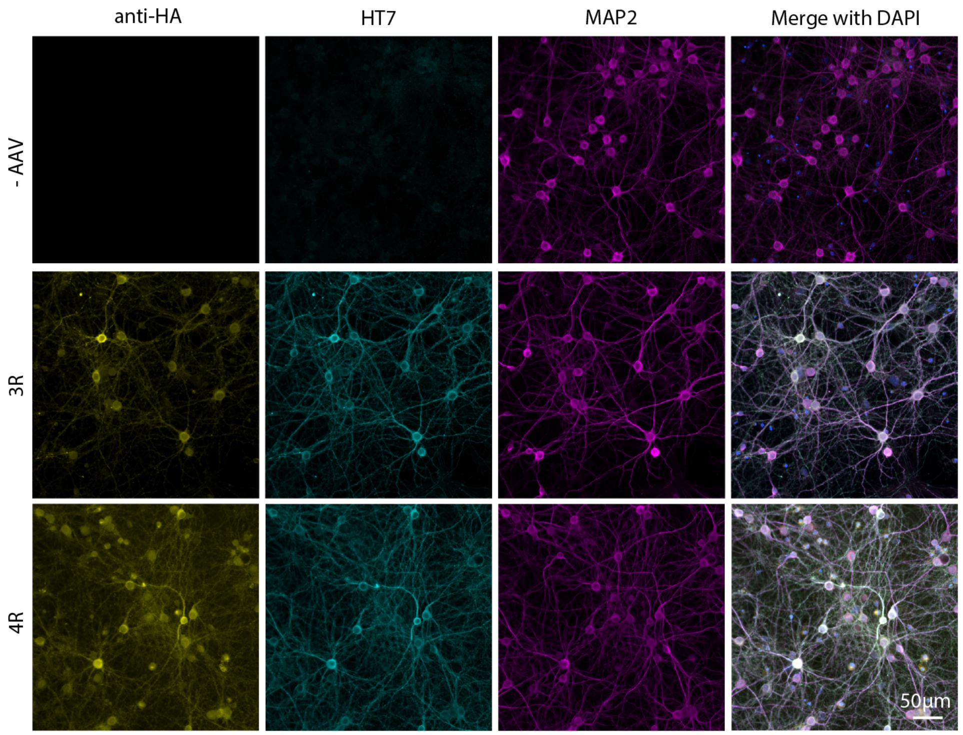


**Figure S1: Expression of human tau isoforms in primary mouse cortical cultures following AAV transduction.**

Immunofluorescence microscopy of primary cultures at DIV14 transduced with AAV1/2-hSyn-HA-0N3R and AAV1/2- hSyn-HA-0N4R, fixed with 4% paraformaldehyde, using anti-HA, anti-tau (HT7), and anti-MAP2 antibodies.

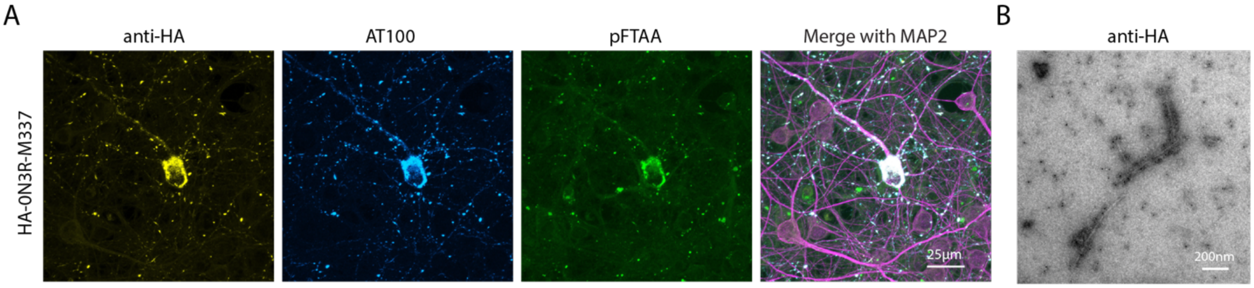


**Figure S2: AD seeded filamentous inclusions form in neurons expressing HA-0N3R-M337 tau.**

A) Representative fluorescence images of primary cultures at DIV21 expressing HA-0N3R-M337 tau and treated with AD seeds. B) Negative stain immunoelectron micrographs of sarkosyl-insoluble extracts using anti-HA antibodies showing gold-labelled filaments.

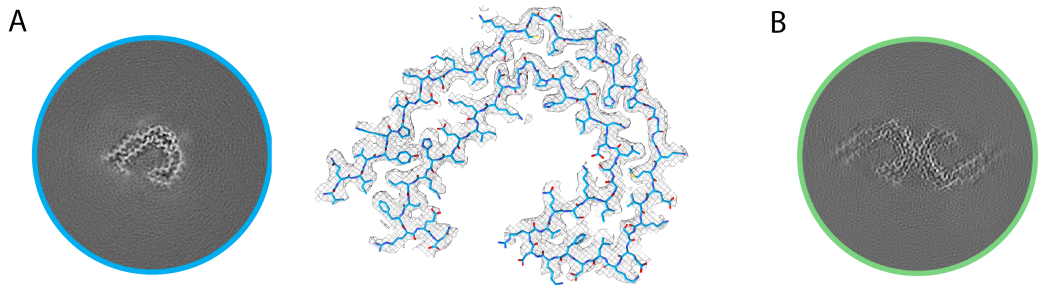


**Figure S3: Cryo-EM density map of single protofilament with the Alzheimer tau fold and cryo-EM density map of PHF from the AD seeds.**

A) Cross-section of reconstructed density of singlet filament extracted from primary cultures expressing HA-0N3R-M337 tau and treated with AD seeds (0.2mg/cm$^2$). Singlet protofilaments with the Alzheimer tau fold represented 18% of filaments and were solved to a resolution of 2.9 Å.
B) Cross-section of reconstructed density of tau filaments extracted from the frontal cortex of AD case 2. The PHF structure was solved to a resolution of 2.7 Å.

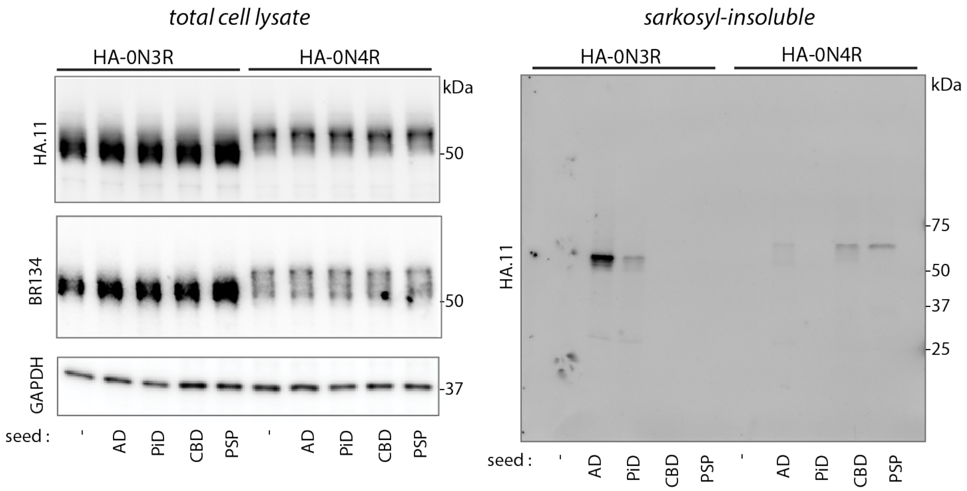


**Figure S4: Seeded aggregation in primary neurons recapitulates disease-specific tau isoform selectivity.**

Immunoblot analysis of total cell lysates and sarkosyl-insoluble fractions of neurons expressing HA-0N3R or HA-0N4R tau and treated with AD, PID, CBD or PSP seeds.

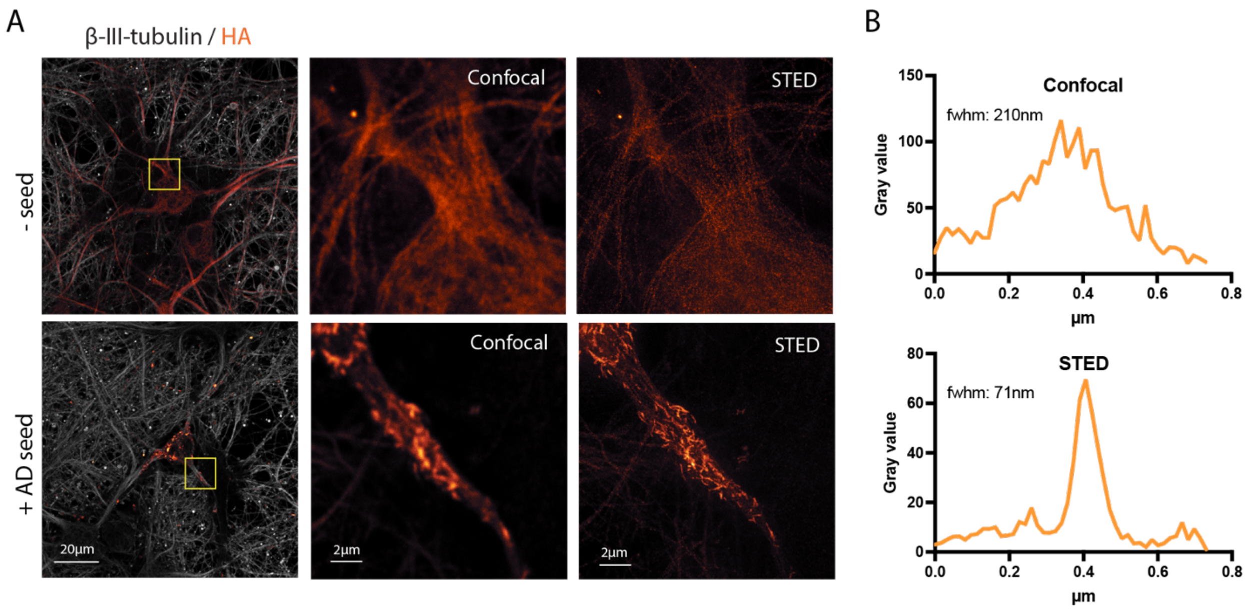


**Figure S5: Super-resolution microscopy imaging of seeded tau inclusions.**

A) Representative confocal and STED images of primary neurons treated +/- AD tau seeds at DIV14 and labelled with β-III-tubulin and anti-HA. B) Representative cross-sectional profile of a filamentous tau inclusion measured by confocal microscopy (top) or STED (bottom).

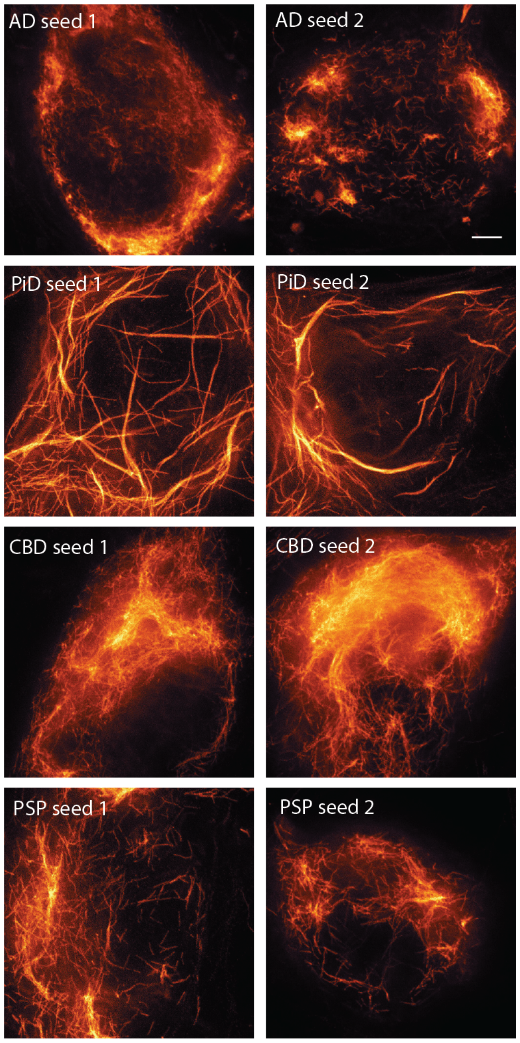


**Figure S6: Tauopathy-derived seeds produce distinct intraneuronal inclusion morphologies.**

STED images at DIV21 of seeded neuronal tau inclusions following the addition of seeds from two cases of AD, PiD, CBD, or PSP. Scale bar, 2μm

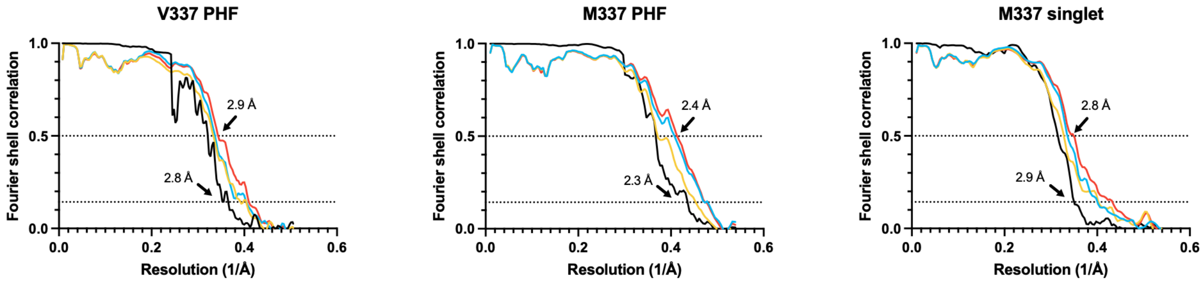


**Figure S7: Resolution estimates for the M337 PHF, M337 singlet, and V337 PHF.** Solvent-corrected Fourier-shell correlation (FSC) curves between independently refined half-maps are shown in black; FSC curves between the refined model and the reconstruction from all particles are shown in red; FSC curves between a model refined against half map 1 against half map 1 are shown in blue; FSC curves of the same model against half map 2 are shown in yellow. Resolution estimates are indicated at thresholds of 0.143 for the half-map FSCs and 0.5 for the model-to-map FSCs

**Table S1. Cryo-EM data acquisition and structure determination**

| | **V-PHF** | **M-PHF** | **M-Singlet** |
|---|---|---|---|
| Data collection | | | |
| Microscope | Titan Krios | | |
| Voltage (kV) | 300 | | |
| Detector | Falcon 4i | | |
| Energy filter slit width (eV) | - | 10 | |
| Magnification | 96,000 | 130,000 | |
| Electron exposure (e-/Å$^2$) | 40 | | |
| Defocus range (μm) | -1.0 to -2.5 | -1.0 to -2.0 | |
| Pixel size (Å) | 0.824* | 0.926 | |
| Data processing | | | |
| Symmetry imposed | C1 | | |
| Initial particle images (no) | 973,577 | 985,972 | 985,972 |
| Final Particle image (no) | 271,595 | 176,278 | 125,248 |
| Map resolution (Å)<br>(FSC threshold 0.143) | 2.8 | 2.3 | 2.9 |
| Helical rise (Å) | 2.41 | 2.43 | 4.88 |
| Helical twist (°) | 179.46 | 179.28 | -1.353 |
| Refinement | | | |
| Initial model | 6HRE | ModelAngelo | |
| Model resolution (Å) FSC threshold | 2.9 | 2.4 | 2.8 |
| Map sharpening *B* factor (Å$^2$) | -56.8 | -53.8 | -51.3 |
| *Model composition* | | | |
| Non-hydrogen atoms | 3528 | 3522 | 1761 |
| Protein residues | 462 | 456 | 228 |
| *B* factors (Å$^2$) | | | |
| Protein | 76.7 | 86.7 | 85.5 |
| *R.m.s. deviations* | | | |
| Bond lengths (Å) | 0.0097 | 0.0101 | 0.0111 |
| Bond angles (°) | 1.80 | 1.75 | 2.19 |
| *Validation* | | | |
| MolProbity score | 1.68 | 1.36 | 1.84 |
| Clashscore | 3.10 | 1.51 | 5.71 |
| Poor rotamers (%) | 0 | 0 | 0 |
| *Ramachandran plot* | | | |
| Favored (%) | 92.00 | 92.57 | 90.54 |
| Allowed (%) | 8.00 | 7.43 | 9.46 |
| Disallowed (%) | 0 | 0 | 0 |

* The V-PHF map was refined at a downscaled pixel size of 0.988 Å.

**Movie S1: Long-term live-cell imaging of progressive tau inclusion formation in primary neurons.** Wide-field images of seeded neurons expressing dendra2-0N3R and seeded at DIV03, taken every 6 h in an Incucyte S3 Live-Cell system.

**Movie S2. High-resolution live-cell imaging of tau inclusions in primary neurons.** Confocal imaging (maximum intensity projections) taken of AD seeded neurons expressing dendra2-0N3R at DIV10 in one field of view over 12 h. Inset: a neuronal cell body with multiple independent events of fluorescent tau inclusion formation.
